# Olfactory receptor 2 drives abdominal aortic aneurysm by promoting CX3CR1-mediated monocyte recruitment

**DOI:** 10.1101/2025.02.28.640923

**Authors:** Patrik Schelemei, Felix Simon Ruben Picard, Yein Park, Philipp Wollnitzke, Harshal Nemade, Dennis Mehrkens, Per Arkenberg, Anna Christine Koebele, Jan Wrobel, Kristel Martinez Lagunas, Elena Wagner, Henning Guthoff, Joy Roy, Moritz Lindquist Liljeqvist, Wiebke Ibing, Markus U. Wagenhäuser, Hubert Schelzig, Bodo Levkau, Ulrich Flögel, Norbert Gerdes, Mohammad Karimpour, Axel M. Hillmer, Gerhard Sengle, Remco T.A. Megens, Christian Weber, Marco Orecchioni, Stephan Baldus, Martin Mollenhauer, Holger Winkels

## Abstract

**Background:** Abdominal aortic aneurysms (AAA) are characterized by an intricate interplay of extracellular matrix degradation and inflammation. Macrophages are centrally involved in these processes. The mechanisms underlying macrophage activation in AAA remain incompletely understood. Vascular macrophages have been shown to express olfactory receptor 2 (Olfr2), a G-protein coupled receptor involved in mediating the sense of smelling and regulating inflammatory activity in macrophages. Whether Olfr2 plays a role in modulating macrophage responses in AAA formation remains unknown.

**Methods & Results:** In silico micro-array analysis showed increased expression of the human Olfr2 orthologue OR6A2 in AAA tissue compared to healthy aorta. Flow cytometric analysis revealed increased expression of OR6A2 on classical, non-classical, and intermediate monocytes of patients with a large AAA (> 5cm) in comparison to patients with smaller AAA (< 5cm). Up to 30% of vascular macrophages expressed OR6A2 in human AAA and Olfr2 in mouse AAA tissue. Olfr2 expression peaked on major histocompatibility complex II-high (MHCII^high^) and C-C chemokine receptor type 2-low (CCR2^low^) aortic monocytes and macrophages on day 7 following experimental AAA initiation and decreased to baseline expression on day 28. Olfr2 gene (*Olfr2^-/-^*) deficiency protected mice from AAA formation, which was accompanied by lowered ECM degradation, reduced macrophage infiltration and increased smooth muscle cell content. Conversely, treatment with the Olfr2 agonist octanal exacerbated AAA formation and inflammation, while the antagonist citral reduced AAA formation in comparison to vehicle treated mice. Bulk transcriptome analysis of aortic tissue revealed reduced inflammatory gene expression in *Olfr2*^-/-^ mice at day 7 following AAA initiation. Spectral flow cytometry resolved 20 aortic immune cell populations, which were largely reduced in quantity by Olfr2-deficiency at day 7 and day 28 post experimental AAA formation, while circulating leukocyte counts and monocyte subset distribution were not altered between *Olfr2*^+/+^ and *Olfr2*^-/-^ mice. Circulating Ly6C^high^-monocytes exhibited reduced expression of the CX3C motif chemokine receptor 1 (CX3CR1) and CCR2 during AAA formation. Transcriptional analysis of monocytes confirmed downregulation of pathways associated with cell adhesion, motility and migration. In vitro, *Olfr2^-/-^* monocytes showed impaired migration towards the CX3CR1 ligand CX3CL1. Competitive transfer of *Olfr2*^+/+^ and *Olfr2*^-/-^ monocytes confirmed reduced migratory capacity of *Olfr2*^-/-^ monocytes into the developing AAA.

**Conclusion:** We demonstrate a critical relevance for Olfr2 in the modulation of the inflammatory response underlying AAA, which is mediated by enhanced monocyte recruitment.

## Introduction

Abdominal aortic aneurysm (AAA) is characterized by progressive structural impairment of the abdominal aorta, resulting in focal arterial enlargement exceeding the normal diameter by more than 50%^1^. The most catastrophic sequela of AAA is acute rupture, which proves fatal in the majority of patients^2, 3^. Next to age and male sex, risk factors for AAA include hypertension, and atherosclerosis^1^. Surgical intervention remains the primary treatment to prevent rupture, while effective pharmacological therapies at early stages are still lacking^4^.

Inflammation is a key feature of AAA formation. Monocyte and macrophage populations have been critically involved in adventitial remodeling and AAA development^5^. Particularly during the early phases of AAA development Ly6C^high^ monocytes infiltrate the aorta and differentiate into macrophages^5^. Monocyte recruitment into the developing AAA has been well-characterized and involves among others the chemokine receptors CCR2, CX3CR1, CXCR4 and CCR5 ^6–9^. Several resident macrophage subtypes reside in the healthy aorta and contribute to homeostatic vessel functions including maintenance of arterial tone, vessel permeability, clearance of apoptotic cells or intraluminal thrombus formation^10, 11^. Recent single cell RNA sequencing studies have uncovered multiple macrophage subsets with inflammatory cytokine (IL-6, TNF-⍺, IL-1β) and extracellular matrix (ECM)-degrading signatures in human^12^ and mouse AAA tissue^13, 14^. Next to inflammation smooth muscle cell (SMC) death and ECM degradation are hallmark features of the pathogenesis underlying AAA. ECM degradation is primarily facilitated by matrix metalloproteinases (MMPs). Macrophages have been shown to secrete MMP9, which has been associated with vessel rupture in patients^15^ and to directly aggravate AAA development in mice^16^. Additionally, secretion of mediators by macrophages such as Netrin-1 has been shown to induce MMP3 activity by surrounding vascular SMCs contributing to AAA progression^17^. While the exact mechanisms governing macrophage activation in AAA are unclear, toll-like receptor signaling, including TLR4, MyD88 and CD14, appear to be central to this process^18^. Recent data demonstrated expression of the olfactory receptor 2 (Olfr2), a G-protein coupled receptor involved in mediating the sense of smelling, on vascular macrophages. Activation of Olfr2 with its homeostatic occurring ligand octanal induced activation of the NLRP3 inflammasome with subsequent IL-1⍺ and IL-1β release in TLR4-primed macrophages^19^. Considering the eminent role of macrophages in AAA and the pivotal function of Olfr2 in macrophage activity, we examined genetic and pharmacological inhibition of Olfr2 in the non-hyperlipidemic porcine pancreas elastase (PPE) infusion AAA mouse model.

## Materials & Methods

Extensive methods are in the Supplemental Material. All raw data and analytical methods are available from the corresponding author on appropriate request.

### Human studies

All clinical samples and measurements were obtained with informed consent from patients after ethics approval (MELENA Study 2018-248-1, biobank 5731R and 2022-2222-TRR259-A09) by the ethics committee of the Heinrich Heine University Düsseldorf. All experiments were performed in accordance with the guidelines of this committee and the Declaration of Helsinki.

### Mouse studies

All animal studies were approved by the local Animal Care and Use Committees [Ministry for Environment, Agriculture, Conservation and Consumer Protection of the State of North Rhine-Westphalia: State Agency for Nature, Environment and Consumer Protection (LANUV), NRW, Germany, AZ: 81-02.04.2021.A106] and conformed to the guidelines from Directive 2010/63/EU of the European Parliament on the protection of animals used for scientific purposes.

### Statistical analysis

Data are presented as mean ± standard deviation. Shapiro–Wilk test and Brown– Forsythe or F-test were used to test for normal distribution and equality of variances, respectively. If data were normally distributed and variances were equal, statistical analysis was performed with Student’s t-test for comparison of two groups. If variances were different Welch correction was applied. For comparison of more than two groups one-way or two-way repeated measures analysis of variance (ANOVA) with post-hoc Tukey’s test was applied. Mann-Whitney (two groups) or Kruskal–Wallis test with Dunn’s post-hoc test (>2 groups) was performed if data were not normally distributed. Fisher’s exact test was used to compare groups of categorical variables. For correlation analysis pearson (normally distributed) or spearman rank test (non-normally distributed) was performed. A p-value <0.05 was considered statistically significant. Possible outliers were assessed by Grubbs (alpha = 0.05). All statistical analyses were performed using GraphPad Prism 8.4.0 (GraphPad Software, San Diego, CA, USA).

## Results

### Olfactory receptor 2 is expressed on macrophages in murine and human aortic aneurysm

To test whether expression of OR6A2, the human Olfr2 ortholog, is altered in human AAA, we performed in-silico micro array analysis of AAA tissue from patients and aortic control tissue^20^. AAA samples were subdivided into media and adventitia and in those covering an intra-luminal thrombus (ILT) or without ILT. OR6A2 expression was significantly increased in media and adventitia of AAA with ILT in comparison to control tissue (**Fig. 1A**). Immunofluorescence staining of human AAA sections (**Sup. Tab. I**) showed OR6A2 expression on CD68^+^ vascular macrophages, of which 30% expressed OR6A2 (**Fig. 1B,C, Sup. Fig 1A)**. To investigate whether OR6A2 is expressed on circulating leukocytes in AAA patients, we performed flow cytometric analysis of PBMCs from age and sex-matched patients categorized by large AAA (5>cm) and small AAA (5<cm). Clinical parameters are listed in **Sup. Tab. II**. In line with previous data^21, 22^, we observed an increase in CD14^+^CD16^+^ intermediate monocytes and NK cells in patients with large AAA (**Fig. 1D, Sup. Fig. 2A,B**). We observed no differences in frequencies of classical and non-classical monocytes, B cells, and multiple T cell populations (**Fig. 1D, Sup. Fig. 2A,B**). While OR6A2 was dimly expressed on B cells, NK cells, and T cells (**Sup. Fig. 2C**), monocytes showed robust OR6A2 expression (**Fig. 1E**). Notably, classical and intermediate monocytes expressed higher levels of OR6A2 in comparison to non-classical monocytes. All three monocyte populations showed higher OR6A2 expression in patients with large AAA compared to small AAA (**Fig. 1E**).

**Figure 1.**
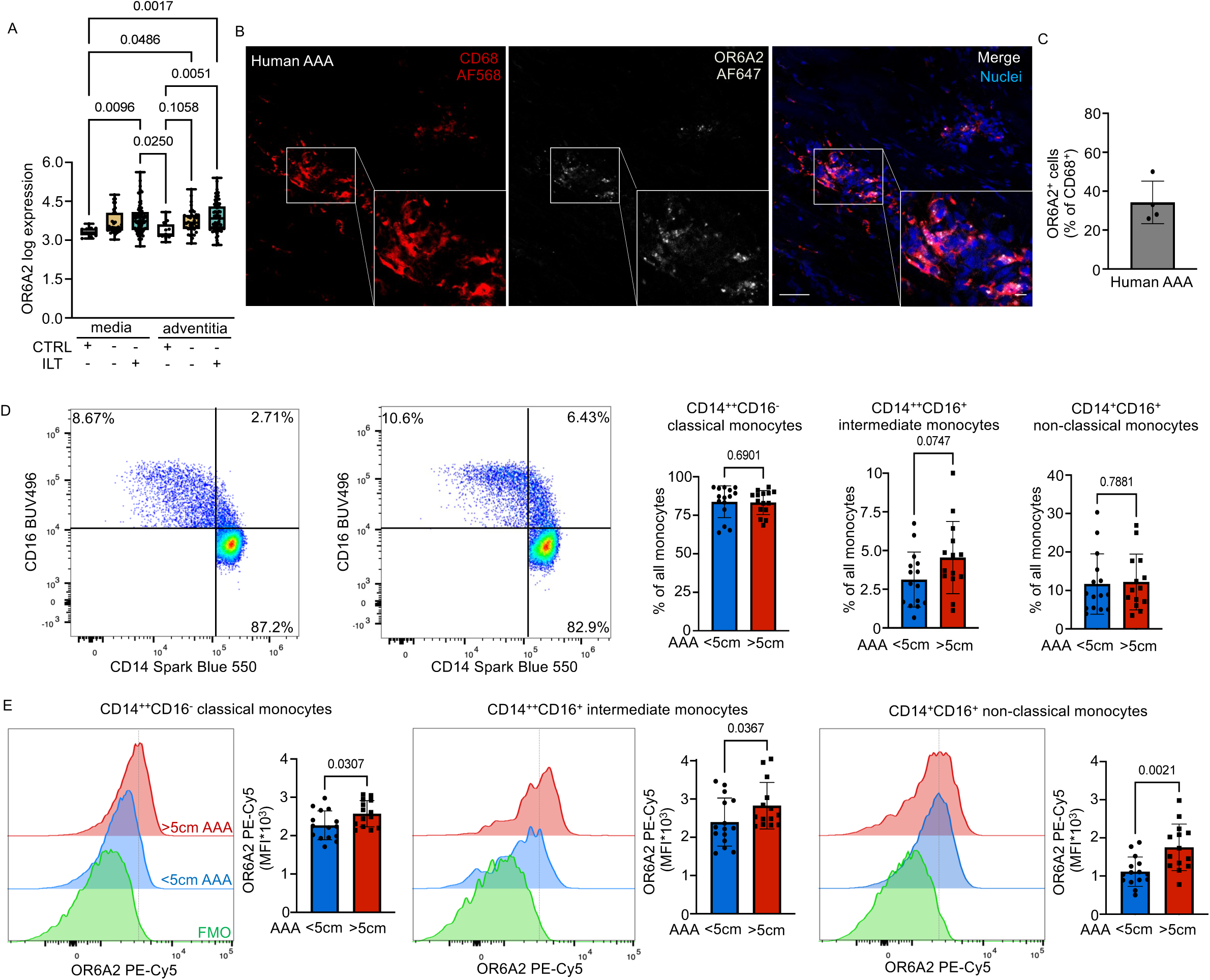
OR6A2 is expressed on circulating monocytes and vascular macrophages in AAA patients. (A) In-silico micro array analysis of OR6A2 expression in human AAA tissue with or without intraluminal thrombus (ILT) and healthy control (CTRL) tissue separated by media and adventitia. Significance was determined by Kruskal-Wallis test with Dunn’s multiple comparisons test. (B) Representative confocal micrograph of human AAA sections stained for OR6A2 (AF647, white), CD68 (AF568, red) and nuclei (Hoechst, blue). Scale bar indicates 50 μm (large panel) and 10 μm (small panel). (C) Quantification of OR6A2-expressing macrophages in human AAA sections. (D) Representative flow cytometry plots and frequencies of monocytes from patients with small (<5cm) and large (>5cm) AAA. (E) Representative histograms and quantification of median fluorescence intensity (MFI) for OR6A2 expression on classical (CD14^++^CD16^-^), intermediate (CD14^++^CD16^+^) and non-classical (CD14^+^CD16^+^) monocytes. Significance was determined by Student’s t-test. Mann-Whitney test was used for non-normally distributed data. Data are presented as mean ± SD.

We induced AAA formation in *Olfr2*^+/+^ and *Olfr2^GFP^* reporter mice by porcine pancreatic elastase (PPE) infusion. Immunofluorescence analysis confirmed a similar frequency of vascular Olfr2-expressing macrophages in aortic sections of *Olfr2^GFP^* reporter mice (**Sup. Fig. 3A-C**) and *Olfr2*^+/+^ mice stained for Olfr2 (**Fig. 2A**). Notably, 90% of all vascular Olfr2-expressing cells were macrophages (**Sup. Fig. 3D**). We next tested by liquid chromatography mass spectrometry/mass spectrometry (LC-MS/MS), whether the Olfr2 agonist octanal might be altered during AAA formation. Plasma octanal levels remained unchanged at 5-6µM at baseline, day 7 and day 28 post experimental AAA formation (**Sup. Fig. 4A,B**). The average aortic concentration was 0.265mg/100g tissue, which is equivalent to 20µM (**Sup. Fig. 4C**). We next aimed to characterize the Olfr2-expressing vascular leukocytes during the peak of the inflammatory (day 7) and reparative/fibrotic phase (day 28). Aortic cells were isolated from *Olfr2*^+/+^ and *Olfr2*^-/-^ mice at day 7 and day 28 and from sham-operated mice. Staining with a 27-marker pan-leukocyte antibody panel resolved 20 populations in an integrated cluster analysis of spectral flow cytometry data comprising: one dendritic cell cluster (cl.10: CD11b^+^CD11c^+^MHCII^+^CD64^-^), two neutrophil cluster (cl.22: CD11b^+^Ly6G^high^CD62L^high^ cl.1: CD11b^+^Ly6G^high^CD62L^low^), three macrophage populations (cl.19: CD11b^+^CD64^+^F4/80^+^CCR2^+^MHCII^+^CX3CR1^+^Ly6C^+^ ; cl.9: CD11b^+^CD64^+^F4/80^+^MHCII^high^CCR2^low^CD11c^+^ ; cl.8: CD11b^+^CD64^+^F4/80^+^TimD4^+^Lyve1^+^CX3CR1^+^CD115^+^), two monocyte cluster (cl.15: CD11b^+^Ly6C^+^CCR2^+^F4/80^low^; cl.7: Ly6C^+^CD16.2^+^MHCII^high^CCR2^int^F4/80^low^) and one myeloid cluster expressing only CD11b and F4/80 (cl.4). Further eight T-cell cluster were detected (CD3^+^): one TCRg/d T-cell population (cl. 18), one effector CD4^+^ T-cell cluster (cl.3: CD62L^-^CD44^+^) and four CD8^+^ T-cell populations. Cytotoxic CD8^+^ T-cells were separated into CD3^+^CD8^+^ (cl.12), CD3^+^CD8^+^Ly6C^+^ (cl.20), CD3^+^CD8^+^CCR2^+^CX3CR1^+^ (cl.13), and CD3^low^CD8^+^CD62L^+^Ly6C^+^ (cl.14) populations. Two T cell populations expressed neither CD4 nor CD8 (cl.17: CD3^+^CD5+; cl.6: CD3^low^CD44+CD69^+^). Two B cell clusters were identified (cl.2: CD19^+^B220^+^MHCII^+^CD62L^+^, cl.23: CD19^+^B220^+^MHCII^+^CD62L^-^). One population expressing only CD62L and CD11c (cl.16) was marked as unknown (**Figure 2C-E**). The cluster analysis uncovered predominant Olfr2 expression by monocyte and macrophage populations (**Fig. 2F**). Two subpopulations modulated Olfr2 expression during the course of AAA formation. MHCII^high^CCR2^low^CD11c^+^ macrophages (cl.9) expressed Olfr2 post sham surgery and significantly upregulated Olfr2 at day 7 post AAA induction, which was reduced to sham levels at day 28. Ly6C^+^CD16.2^+^MHCII^high^CCR2^int^ monocytes (cl.7) did not express Olfr2 upon sham surgery, while Olfr2 expression peaked at day 7 and decreased at day 28 (**Fig. 2G**). MHCII^int^CCR2^+^ macrophages (cl.19) and Ly6C^+^CCR2^high^ monocytes (cl.15) expressed similar Olfr2 levels after sham surgery and at days 7 and 28 of AAA formation. Resident-like macrophages (cl.8) and the CD11b^+^F4/80^+^ population (cl.4) did not express Olfr2 **(Sup. Fig 5A)**. Of note, circulating Ly6C^high^ monocytes showed significant Olfr2 upregulation at day 7 post PPE, which remained elevated at day 28, while Ly6C^low^ monocytes did not significantly regulate Olfr2 expression during AAA formation (**Sup. Fig 6B).**

**Figure 2.**
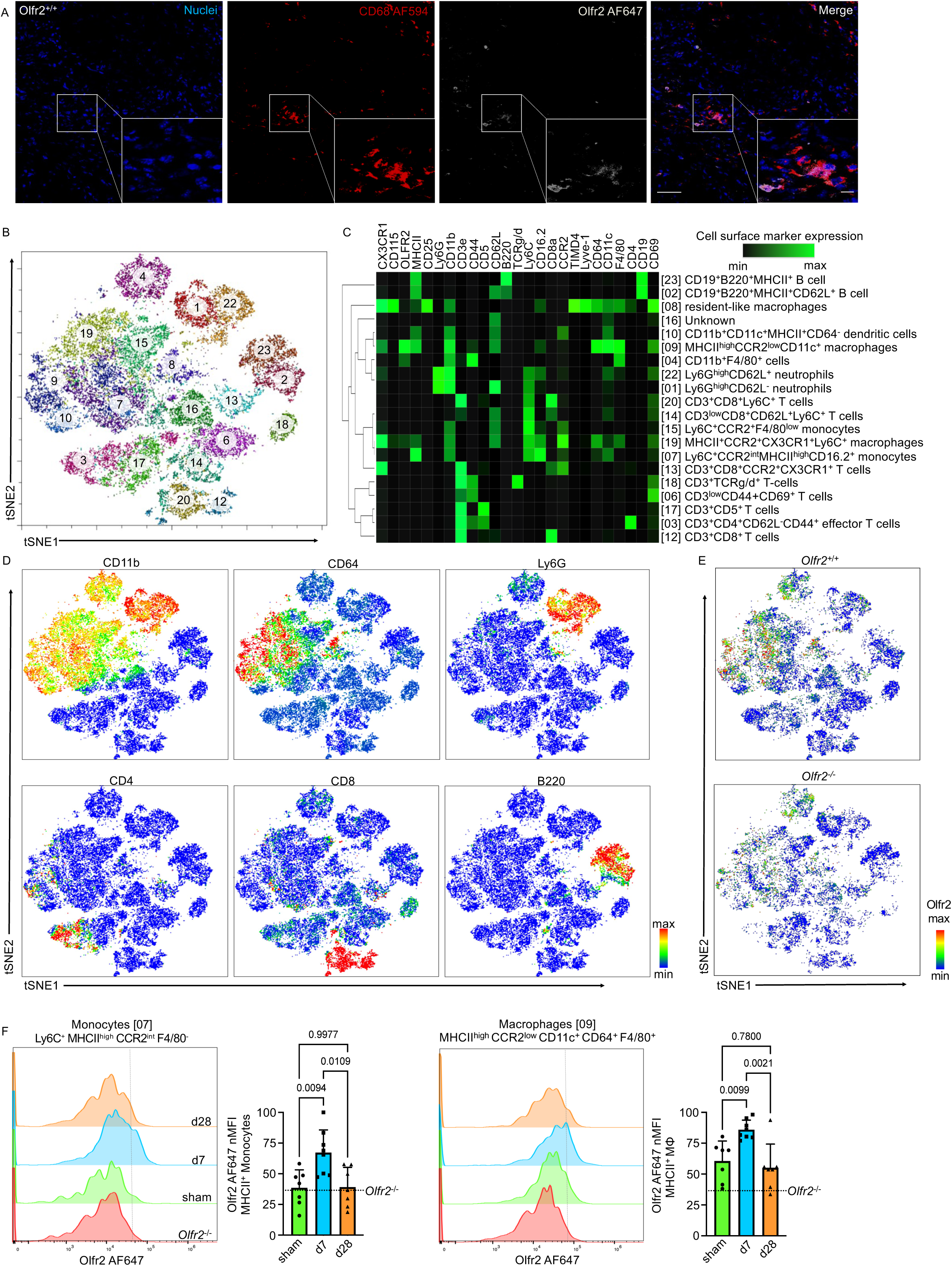
Olfr2 is dynamically expressed by mouse monocytes and macrophages. (A) AAA was induced in *Olfr2*^+/+^ by porcine pancreatic elastase infusion into the infrarenal aorta. Representative confocal micrograph showing Olfr2 (AF647, white), CD68 (AF594, red) and nuclei (Hoechst, blue) immunofluorescence staining in a whole-mount infrarenal aorta of *Olfr2*^+/+^ mice. Scale bar indicates 50 μm (large panel) and 10 μm (small panel). (B) Unsupervised cell cluster detection by t-distributed stochastic neighbor embedding (tSNE) and cluster detection algorithm (PhenoGraph) from spectral flow cytometry (FACS) analysis of aortic leukocytes at day 7 and day 28 after PPE and sham surgery (pooled). (C) Hierarchically clustered heatmap of column normalized FACS marker expression levels in the annotated leukocyte subtypes derived from clustering (right). (D) Expression of CD11b, CD64, Ly6G, CD4, CD8 and B220 projected on the tSNE map. (E) Projection of Olfr2 onto tSNE maps of aortic leukocytes from *Olfr2*^+/+^ and *Olfr2*^-/-^ mice. (F) Representative histograms and flow cytometric analysis of normalized median fluorescence intensity (nMFI) of Olfr2 expression on Ly6C^+^MHCII^+^ monocytes [cl. 7] and MHCII^high^CCR2^low^ macrophages [cl. 9] in the infrarenal aorta at day 7 and day 28 after PPE as well as sham surgery. Dotted line indicates nMFI of the respective *Olfr2*^-/-^ cluster. Significance was determined by ordinary one-way ANOVA with Tukey’s multiple comparisons test. Data are presented as mean ± SD.

Taken together, this data indicates that circulating Ly6C^high^ monocytes, vascular MHC^+^CCR2^low^ monocytes and macrophages dynamically regulate Olfr2 expression during AAA formation.

### Olfr2-deficiency reduces AAA formation

To uncover whether Olfr2 plays a role in AAA formation, *Olfr2^-/-^* mice and littermate controls (*Olfr2^+/+^*) underwent PPE infusion. Ultrasound studies demonstrated a two-fold enlarged aortic diameter relative to baseline in *Olfr2^+/+^* mice 28 days following AAA induction, while *Olfr2^-/-^* mice were protected from AAA formation (**Fig. 3A,B**). As ECM degradation is a hallmark of AAA formation, we determined whether lack of Olfr2 might also protect from AAA-associated ultrastructural changes of the aortic wall. Histologically, *Olfr2^-/-^*mice exhibited preserved elastin structure while *Olfr2^+/+^* mice showed signs of elastin degradation characterized by fiber thinning, fragmentation, and medial degeneration (**Fig 3C**). Second harmonics generation (SHG) in two-photon-microscopy showed increased collagen deposition (**Fig. 3D**) in *Olfr2^-/-^* mice, that was corroborated by preserved SMC content (**Fig. 3E**). Lastly, in comparison to *Olfr2^+/+^* mice, AAA tissue from *Olfr2^-/-^* mice exhibited less macrophages assayed by CD68 staining (**Fig. 3F**). These data show that Olfr2 affects AAA development by aggravating pathological ECM remodeling and macrophage infiltration.

**Figure 3.**
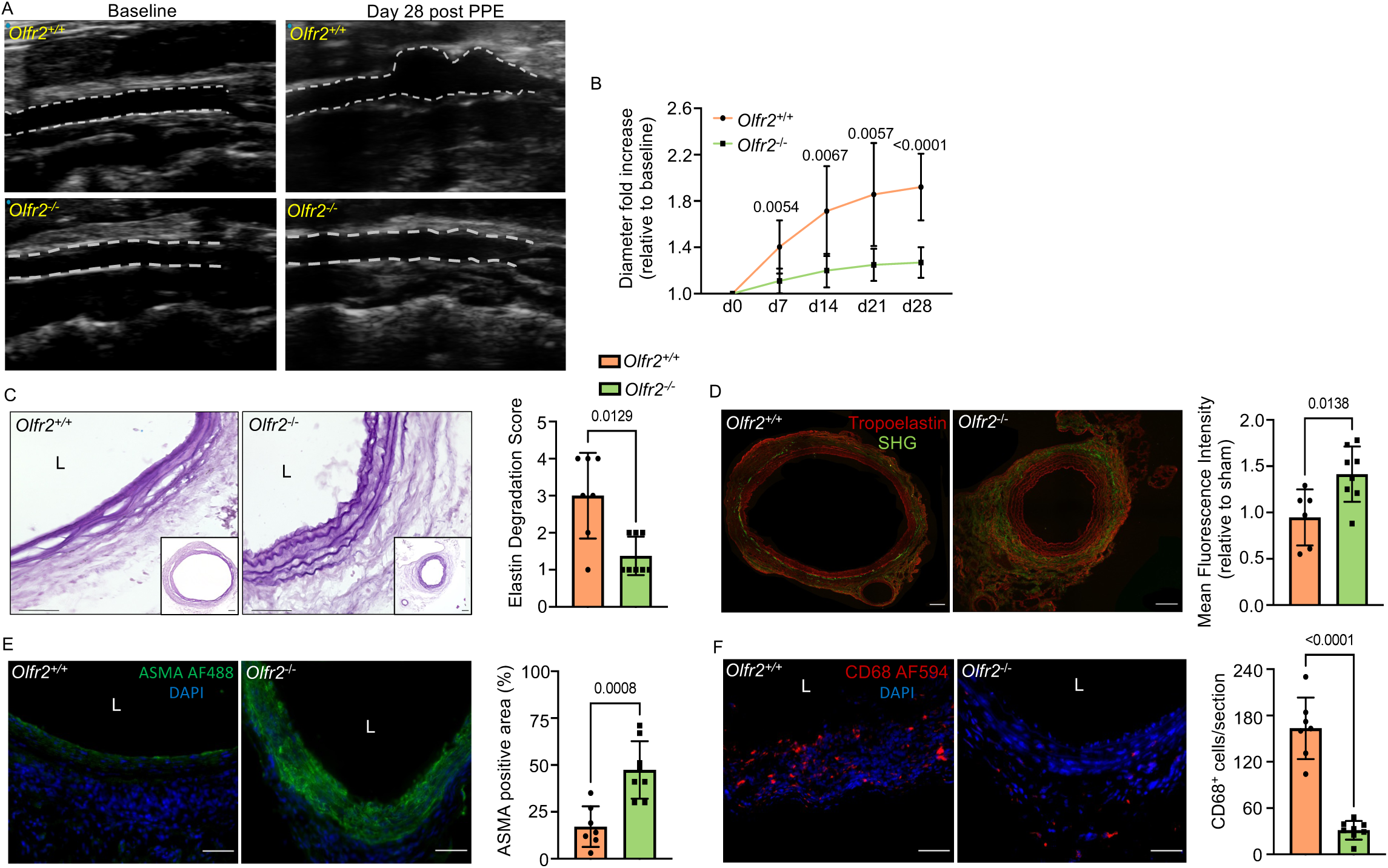
Olfr2-deficiency reduces AAA formation. (A) Representative ultrasound images of aortas of *Olfr2*^+/+^ (WT) and *Olfr2*^-/-^ mice at baseline and day 28 post PPE. (B) Analysis of the aortic diameter of WT and Olfr2^-/-^ mice at baseline (d0), day 7, day 14, day 21 and day 28 after aneurysm induction. Data is visualized as the diameter fold change relative to the aortic diameter at baseline (n=8 per group). Significance was determined by Student’s t-test. Welch correction was applied if variance was unequal. (C) Representative images of histological elastica staining. Quantification of elastin degradation indices have been performed for each group. L = Lumen. Scale bar indicates 50 μm (large panel) and 100 μm (small panel). Significance was determined by Mann-Whitney test. (D) Representative images of tropoelastin immunofluorescence staining (red) and collagen visualized by second harmonic generation (green) in AAA sections. Collagen levels were analyzed as mean fluorescence intensity of the SHG signal. Scale bar indicates 100 μm. Significance was determined by Student’s t-test. (E) Representative alpha smooth muscle cell actin (ASMA) immunofluorescence staining in AAA sections and quantification of the positive area as percentage of the aortic media. Nuclei are stained with DAPI (blue) Scale bar indicates 50 μm. Significance was determined by Student’s t-test. (F) Representative immunofluorescence images of AAA cross-sections stained for CD68 (red) with quantification of CD68^+^ cells in the whole section. Nuclei are stained with DAPI (blue). Scale bar indicates 50 μm. Significance was determined by Student’s t-test with Welch correction. Data are presented as mean ± SD.

### Pharmacological modulation of Olfr2 affects AAA formation

To test whether pharmacological modulation of Olfr2 affects AAA formation, we injected C57BL6/J mice intraperitoneally with octanal or the Olfr2 inhibitor citral^23^, both at 10 μg per gram body weight, every three days during the 28-day observation period post PPE infusion. (**Fig. 4A**). Octanal treatment significantly increased the aortic diameter in comparison to vehicle controls, while citral treatment reduced the aortic diameter (**Fig. 4B-C**). Notably, Olfr2-deficient mice treated with vehicle or octanal developed similar aneurysm sizes as achieved by citral injections. Octanal treatment increased infiltration of aortic macrophages (**Fig. 4F**). In contrast, citral treatment preserved SMC content and decreased vascular macrophage content (**Fig. 4E,F**). Although qualitative differences in elastin degradation of citral (decreased) and octanal (increased) treated mice compared to vehicle controls were observed, they were not statistically significant (**Fig. 4D**).

**Figure 4.**
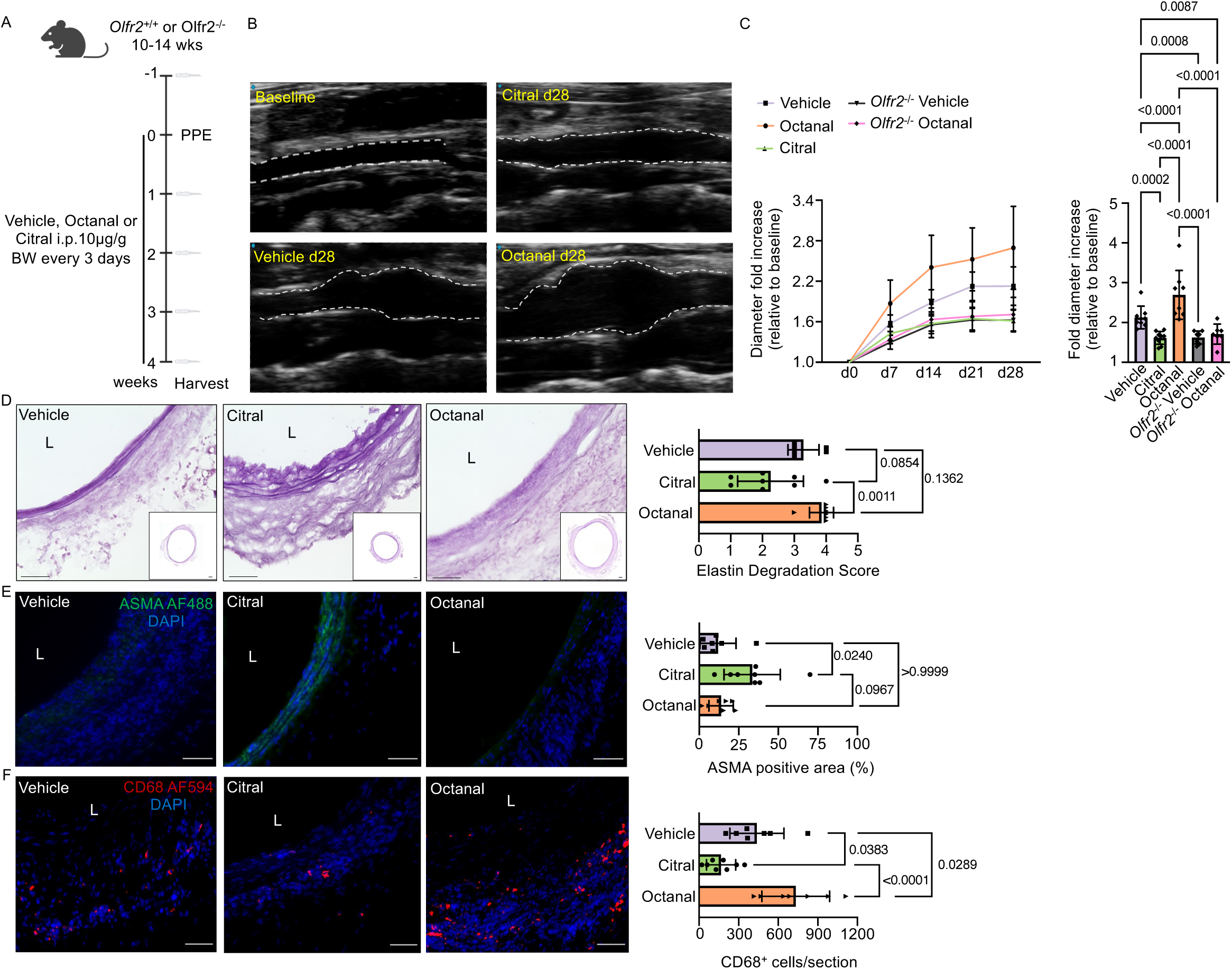
Pharmacological Olfr2 inhibition reduces AAA formation. (A) Treatment protocol for (B-F). BW = body weight. (B) Representative ultrasound images of aortas of WT mice treated with octanal, citral or vehicle at baseline and day 28 post PPE. (C) Analysis of the aortic diameter of the indicated groups at baseline (d0), day 7, day 14, day 21 and day 28 after aneurysm induction. Data is visualized as the diameter fold change relative to the aortic diameter at baseline. P-values are only shown for day 28 in the bar graph (right panel) for clearer visibility. Significance was determined by ordinary two-way ANOVA with Tukey’s multiple comparisons test. (D) Representative images of histological elastica staining. Quantification of elastin degradation indices have been performed for each group. L = Lumen. Scale bar indicates 50 μm (large panel) and 100 μm (small panel). Significance was determined by Kruskal-Wallis test with Dunn’s multiple comparisons test. (E) Representative alpha smooth muscle cell actin (ASMA) immunofluorescence staining in AAA sections and quantification of the positive area as percentage of the aortic media. Nuclei are stained with DAPI (blue). Scale bar indicates 50 μm. Significance was determined by Kruskal-Wallis test with Dunn’s multiple comparisons test. (F) Representative immunofluorescence images of AAA cross-sections stained for CD68 (red) with quantification of CD68^+^ cells in the whole section. Nuclei are stained with DAPI (blue). Scale bar indicates 50 μm. Significance was determined by ordinary one-way ANOVA with Tukey’s multiple comparisons test. Data are presented as mean ± SD.

### Olfr2-deletion reduces vascular inflammation in AAA

Bulk transcriptome analysis of infrarenal AAA tissue collected at day 7 from *Olfr2^+/+^* and *Olfr2^-/-^* mice was performed to uncover molecular mechanisms underlying the protective action of Olfr2 inactivation in AAA. In total, 356 genes were differently expressed (DE), of which 166 were up-and 290 were downregulated when Olfr2 was absent (**Fig. 5A, Sup. Fig.6A, Sup. Tab. V**). Gene ontology analysis of DE-genes showed enrichment of immune and inflammation pathways among the top 20 pathways in AAA tissue of *Olfr2^+/+^* mice (**Fig. 5B**). In contrast, AAA tissue from *Olfr2^-/-^*mice was enriched for multiple gene terms related to vascular SMCs, ECM organization, and energy metabolism (**Sup. Fig. 6B)**. These findings are in line with our histological analysis and prompted us to investigate the immune response in more detail. Systemically, Olfr2-deficiency significantly reduced circulating levels of IL-5 at day 7 whereas TNF⍺, IL-6, IL-9 and IL-10 were significantly reduced at day 28. **(Sup. Fig. 7**). Spectral flow cytometric analysis of digested AAA tissue revealed a ten-fold increase in leukocyte counts comprising multiple myeloid and lymphoid populations at day 7 in comparison to day 28 of AAA in both, *Olfr2^+/+^* and *Olfr2^-/-^* mice, highlighting the peak of the inflammatory response in AAA at day 7 (**Fig. 5C-D**). Notably, genetic Olfr2-deficiency reduced aortic leukocyte counts profoundly at both timepoints compared to *Olfr2*^+/+^ (**Fig. 5D**). This was particularly driven by a reduction Ly6C^+^MHCII^+^ (cl. 7) monocytes MHCII^+^CCR2^low^ (cl. 9) macrophages, which demonstrated dynamic Olfr2 expression, and MHCII^int^CCR2^+^ (cl. 19) macrophages at day 7 and day 28 of AAA formation, respectively (**Fig. 5E**). Of note, AAA induction did not alter circulating leukocyte counts or the distribution of Ly6C^high^ and Ly6C^low^ monocytes between *Olfr2^+/+^* and *Olfr2^-/-^* mice at day 7 and day 28 (**Sup. Fig. 8)**.

**Figure 5.**
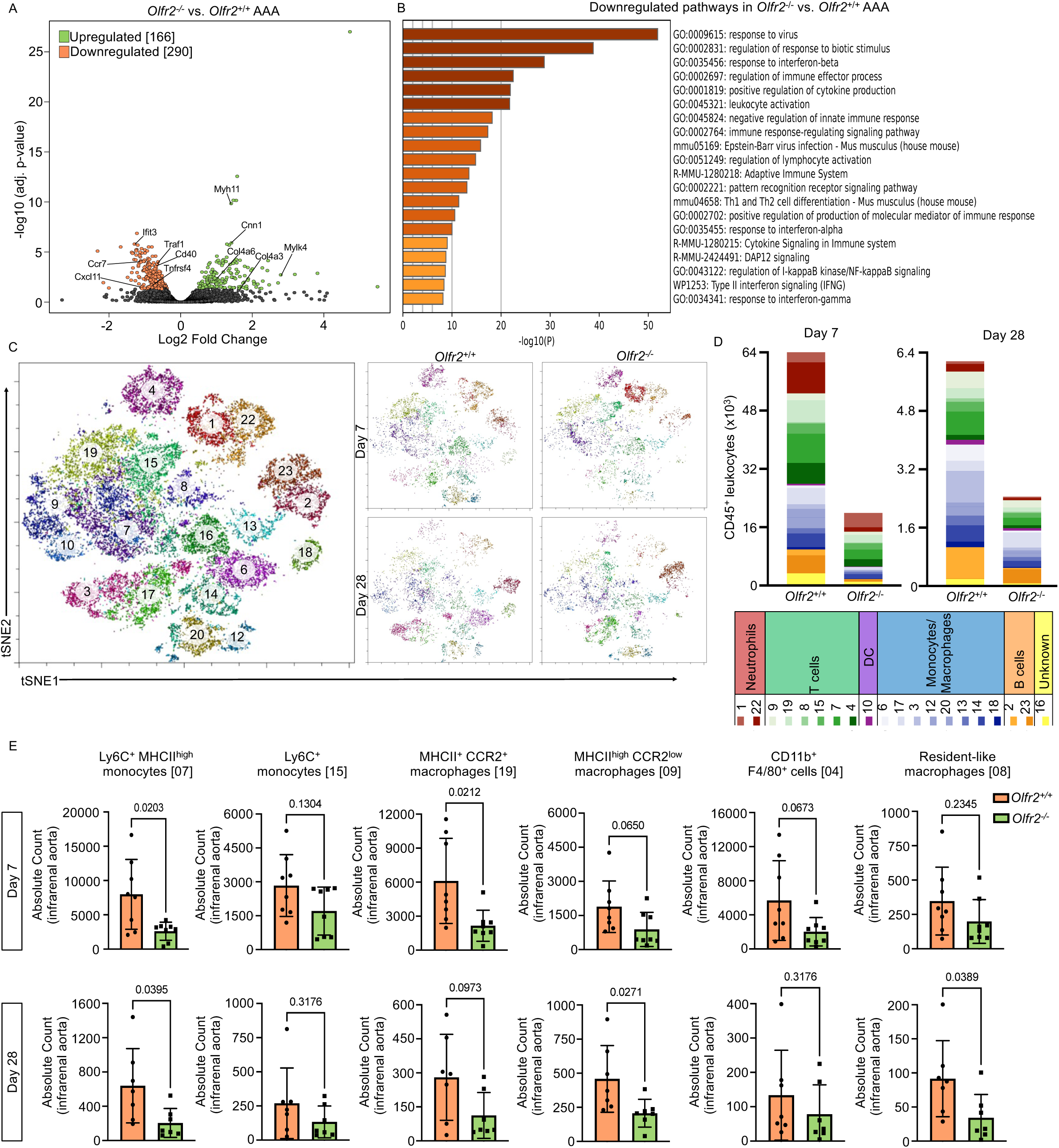
Olfr2-deletion reduces vascular inflammation in AAA. (A) Volcano plot of bulk RNA sequencing of *Olfr2*^-/-^ vs. *Olfr2*^+/+^ AAA tissue at day 7 post PPE. (B) Top 20 downregulated pathways in *Olfr2*^-/-^ vs *Olfr2*^+/+^ AAA generated by Metascape. (C) Unsupervised cell cluster detection by tSNE and PhenoGraph from spectral flow cytometry (FACS) of aortic leukocytes (pooled; left panel) and tSNEs separated by genotype and time point post PPE (day 7 and day 28; right panel). (D) Bar chart of the mean count of immune cell types from *Olfr2*^+/+^ and *Olfr2*^-/-^ AAA tissue at day 7 and day 28 post PPE derived from flow cytometry data. (E) Quantification of monocyte and macrophage subclusters at day 7 and day 28 post PPE in *Olfr2*^+/+^ and *Olfr2*^-/-^ AAA tissue. Significance was determined by Student’s t*-*test. Welch correction was applied if variance was unequal. Mann-Whitney test was used for non-normally distributed data. Data are presented as mean ± SD.

### Olfr2 promotes migration of Ly6C^high^ monocytes into AAA

Ly6C^high^ monocytes are progenitors of AAA-associated macrophages. The equal numbers of circulating monocytes between *Olfr2^+/+^* and *Olfr2^-/-^* mice in AAA formation yet reduced monocyte and macrophage accumulation in *Olfr2*^-/-^ mice prompted us to investigate these cells further. Ly6C^high^ monocytes were isolated from *Olfr2^+/+^*and *Olfr2^-/-^* mice at day 7 post AAA induction for transcriptional profiling. In total, 755 DE genes were detected of which 325 genes were up-and 430 downregulated (**Fig. 6A, Sup. Fig. 9A, Sup. Tab. VI)**. Upregulated genes were mostly associated with cell cycle terms (**Fig. 6B, Sup. Fig. 9B**). Pathway analysis of downregulated genes in *Olfr2^-/-^*monocytes showed attenuation of an inflammatory response which included significantly downregulated expression of *Relb*, *Junb*, and *Ncoa6* (**Sup. Tab. VI**). Gene set enrichment analysis of in *Olfr2^-/-^* monocytes significantly DE genes showed reduced enrichment of TNF⍺ signaling via NFκB and cell activation, indicating reduced inflammatory activation (**Sup. Fig. 9C**). Furthermore, GO terms related to myeloid cell differentiation were downregulated (**Fig. 6B**). This was reflected by low expression of several key transcription factors involved in monocyte differentiation and maturation including *Nr4a1*, *Cebpb, Socs1,* and *Irf7*^24, 25^ in *Olfr2^-/-^* monocytes (**Fig. 6C**). *Olfr2^-/-^*monocytes furthermore showed profound downregulation of aggregated pathways regulating cell adhesion and cell motility (**Fig. 6B**), which included cell migration (GO:0030335; p=0.0004). Attenuated monocyte differentiation has been linked to lower expression of the chemokine receptors CCR2 and CX3CR1^26^. FACS analysis of Ly6C^high^ monocytes from *Olfr2^-/-^* and *Olfr2*^+/+^ mice revealed decreased expression of the chemokine receptors CX3CR1 and CCR2 on *Olfr2^-/-^* monocytes in comparison to *Olfr2*^+/+^ at day 7 of AAA formation, which returned to baseline at day 28. Notably, CX3CR1 expression remained lower on *Olfr2^-/-^*monocytes at day 28 post-AAA induction (**Fig. 6D**). Interestingly, expression of OR6A2 positively correlated with increased CX3CR1 and CCR2 expression on circulating classical and non-classical monocytes in AAA patients (**Sup. Fig. 10)**. To test whether the reduced CX3CR1 expression on *Olfr2^-/-^*monocytes is functionally relevant, monocytes were isolated from *Olfr2^+/+^*and *Olfr2^-/-^* mice and subjected to in vitro migration using transwell migration plates with 5μm pore size. Chemotaxis of Olfr2-deficient monocytes to the CX3CR1 ligand CX3CL1 was significantly reduced (**Fig. 6E**). As CX3CR1 has been implicated in monocyte recruitment and adhesion^27, 28^, we examined adhesion of Olfr2-deficient monocytes to the vascular wall in an ex vivo perfusion assay. Carotid arteries from C57BL6/J were stimulated for 3 h with TNF⍺, mounted in a perfusion chamber, and perfused with Green CMFDA-and Deep Red cell tracker-stained monocytes isolated from *Olfr2^+/+^* and *Olfr2^-/-^* mice. Multi-photon microscopy of mounted carotids uncovered that significantly less *Olfr2^-/-^*monocytes adhered to the endothelium in comparison to *Olfr2^+/+^*monocytes (**Fig. 6F**). To assess whether the reduced in vitro migration and ex vivo adhesion might translate into reduced monocyte recruitment in vivo in AAA, monocytes were isolated from *Olfr2^+/+^* and *Olfr2^-/-^* mice, stained with cell trace violet and cell trace yellow, respectively, and intravenously injected into C57BL6/J mice at day 3 of AAA formation. Flow cytometric analysis of intrarenal AAA tissue 24 hours post adoptive transfer demonstrated reduced accumulation of *Olfr2^-/-^* monocytes (**Fig. 6G,H, Sup. Fig. 11**). Collectively, this data suggests that Olfr2-deficiency causes monocyte intrinsic alterations that lead to impaired recruitment into the aortic wall, predominantly driven by reduced CX3CR1 expression.

**Figure 6.**
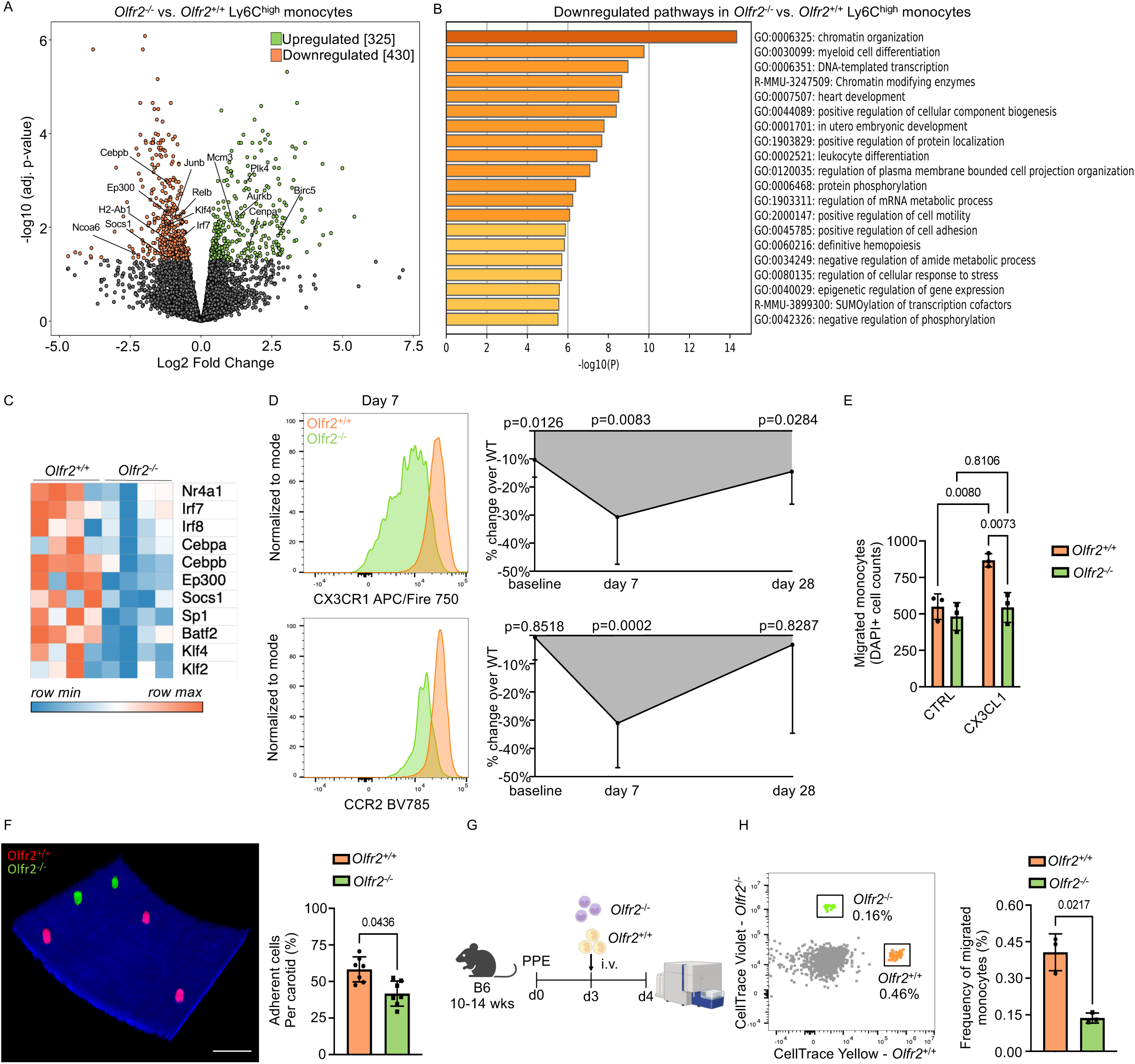
Olfr2 promotes migration of Ly6C^high^ monocytes into AAA. (A) Volcano plot of bulk RNA sequencing of splenic Ly6C^high^ monocytes of *Olfr2*^-/-^ vs. *Olfr2*^+/+^ mice at day 7 post PPE. (B) Top 20 downregulated pathways in *Olfr2*^-/-^ vs *Olfr2*^+/+^ Ly6C^high^ monocytes. (C) Heatmap of row-normalized expression of key transcription factors involved in myeloid cell differentiation. (D) Representative histograms and quantification of CX3CR1 and CCR2 expression on Ly6C^high^ monocytes by median fluorescence intensity at indicated time points after PPE. Results are shown as mean ± SD percent change of expression in *Olfr2*^-/-^ compared to *Olfr2*^+/+^. n=7-8 per group and time point. Significance was determined by Student’s t-test. (E) In vitro chemotaxis of isolated monocytes towards CX3CL1 (25ng/ml) in Boyden chambers. Migrated cells at the underside of the membrane were counted in 5 random fields. Significance was determined by ordinary two-way ANOVA with Tukey’s multiple comparisons test. (F) Representative image and quantification of *Olfr2*^+/+^ (red) and *Olfr2*^-/-^ (green) monocytes adhering to TNF-activated endothelium in an ex-vivo perfusion assay. Scale bar indicates 50 μm. Significance was determined by paired *t* test. (G) Experimental outline for adoptive transfer of labeled *Olfr2*^+/+^ (CellTrace yellow) and *Olfr2*^-/-^ (CellTrace violet) monocytes (1 x 10^6^ each) into C57BL6/J mice with a developing AAA at day 3 post PPE. Aortic tissue was harvested for flow cytometry 24 after transfer. (H) Representative flow cytometry plot and frequencies of adoptively transferred CellTrace^+^ *Olfr2*^+/+^ and *Olfr2*^-/-^ cells in recipient aortic tissue 24h post adoptive transfer. Significance was determined by paired *t* test. Data are presented as mean ± SD.

## Discussion

We here report a crucial role for Olfr2 in AAA development. Patients with AAA >5cm exhibited higher OR6A2 expression on circulating monocytes in comparison to patients with smaller AAAs, which emphasizes the potential disease-and stage-specific relevance of this pathway. OR6A2/Olfr2 expression was identified almost exclusively on monocytes and macrophages in human and murine AAA tissue. Genetic Olfr2 deficiency and pharmacological Olfr2 inhibition significantly reduced AAA formation in mice. We highlight a novel mechanism by which deficiency of Olfr2 reduced expression of CX3CR1 on classical monocytes and impairs their recruitment to the AAA wall, which further abrogates the subsequent inflammatory response required for the development of AAA.

Infiltration of myeloid cells and subsequent macrophage activation is a key hallmark of AAA and is linked to extensive pathological remodeling of the ECM^5^. Cytokines such as IL-1β and TNF-⍺ have been shown to induce inflammatory macrophage polarization, SMC phenotype switch or even apoptosis^5^. Depletion of monocytes and macrophages by clodronate liposomes^29^ or in genetic models^30^ has been shown to ameliorate aortic aneurysm formation in mouse models.

A recent study showed expression of Olfr2 on vascular macrophages while its activation induced NLRP3-dependent IL-1β and TNF⍺ secretion and Olfr2-deficiency ameliorated atherosclerosis^19^. In line with these findings, we here show significantly reduced AAA formation in *Olfr2*^-/-^ mice, which was driven by systemic and local attenuation of inflammation. *Olfr2*^-/-^ mice had decreased numbers of vascular macrophages and classical monocytes in AAA tissue during the peak of the inflammatory (d 7) and the remodeling phase (d 28). The accumulation of AAA-associated macrophages is driven by recruitment of Ly6C^high^CX3CR1^+^ classical monocytes, which are pro-inflammatory and differentiate into macrophages within tissue following extravasation^26, 31^. Indeed, expression of CX3CR1 positively correlated with increased inflammatory macrophage content in human AAA^9^.

The extravasation of monocytes into inflamed tissue involves a well-orchestrated cascade^32^, which among others depends on selectins, GM-CSF^33^, Trem1^34^, and the chemokine receptors CCR2, CX3CR1, CCR5 as well as their respective ligands, CCL2, CX3CL1 and CCL5/CCR5^5, 35^. Deficiency or blockade of CCR2 or CX3CR1 and their ligands, has been shown to protect against experimental AAA formation^36, 37^ ^38^. We here demonstrate reduced CCR2 and CX3CR1 expression on circulating Ly6C^high^ monocytes at day 7 of aneurysm formation in *Olfr2*^-/-^ mice suggesting impaired inflammatory signaling in the absence of Olfr2. CX3CR1 expression remained lower on Ly6C^high^ monocytes from *Olfr2*^-/-^ mice in comparison to controls at day 28 post AAA induction, while CCR2 returned to baseline levels. Notably, OR6A2 expression correlated with increased CX3CR1 and CCR2 on human monocytes, suggesting that OR6A2 might directly or indirectly act on expression of these receptors also in humans. Several factors, such as TGFβ^39^, IL-10^40^, IFNγ^40^ and angiotensin-II^41^ have been identified to upregulate or stabilize CX3CR1 expression on monocytes. Of those IFNγ and IL-10 were included in the plasma cytokine analysis of *Olfr2^+/+^* and *Olfr2*^-/-^ mice. While IFNγ was barely detected in both genotypes, IL-10 levels were decreased at day 28 in *Olfr2*^-/-^ mice in comparison to *Olfr2^+/+^* mice. The reduced abundance of inflammatory cytokines might contribute to decreased CX3CR1 levels on Ly6C^high^ monocytes in *Olfr2*^-/-^ mice. Monocytes from *Olfr2^-/-^* mice further exhibited reduced expression of genes involved in cell adhesion in and impaired adhesion to inflamed endothelium, which might further contribute to the observed reduction in monocytes and macrophages in AAA of *Olfr2*^-/-^ mice. Next to reduced monocyte migration, we observed an effect of Olfr2-deficiency on monocyte function, which as well might contribute to reduced AAA formation. While steady Olfr2 expression was detected on vascular CCR2^+^Ly6C^+^ monocytes and early macrophages characterized by Ly6C, CCR2 and CX3CR1 expression^26^, Olfr2 was only increased on MHCII^high^ monocytes and macrophages during the inflammatory phase of AAA. Previous work showed that Ly6C^+^F4/80^-^ monocytes recruited to the inflamed colon mature by first acquiring MHCII, followed by F4/80 expression upon which Ly6C expression is lost^42, 43^. This maturation coincided with increased inflammatory gene expression^42, 43^. While the macrophage clusters identified by spectral flow cytometry here warrant further investigation, the MHCII expression on monocytes and macrophages dynamically regulating Olfr2 expression in the course of AAA suggests a more inflammatory, activated and mature phenotype, respectively. This is in line with a previous publication identifying Olfr2-expressing monocytes and macrophages as pro-inflammatory^44^. We here show attenuated inflammation in *Olfr2*^-/-^ monocytes evidenced by reduced expression of *Relb, Ncoa6,* and *Junb*, all mediators that are critically involved in NFκB signaling and inflammatory cytokine production^45–47^. Moreover, multiple transcription factors and mediators of monocyte differentiation including *Nr4a1*, *Cebpb, Socs1,* and *Irf7,* were decreased in *Olfr2*^-/-^ monocytes. These four genes are lowly expressed by myeloid progenitor cells in the bone marrow and increase in differentiated circulating monocytes^24^. *Cebpb*, which encodes for (CCAAT/enhancer-binding protein beta) C/EBPβ, can bind the Nr4a1 promotor for Ly6C^low^ monocyte generation,^24, 48^. While the role for Nr4a1 in Ly6C^low^ monocyte generation is well established, competitive bone marrow transplantation also resulted in reduced Ly6C^high^ monocytes derived from *Nr4a1*^-/-^ bone marrow compared to WT subsets, indicating a broader role for Nr4a1 in monocyte development.

Suppressor of cytokine signalling 1 (SOCS1) has been shown to engage in a transcription factor signaling network including *Pparg*, *Nr4a1* and *Cx3cr1*, which are important for monocyte differentiation and survival^49, 50^. Interestingly, genetic deficiency of *Socs1* lead to a reduction in CX3CR1 expression in both Ly6C^high^ and Ly6C^low^ monocytes, though this effect appeared to be stronger in Ly6C^high^ monocytes^51^. Notably, circulating leukocytes and the ratio of Ly6C^high^ to Ly6C^low^ monocytes were not affected by Olfr2-deficiency, suggesting that the effect of Olfr2-deficiency alters rather the functionality of mature monocytes than their development per se. Interestingly, while we observed a reduction in infiltrating macrophages at the aneurysm site in *Olfr2*^-/-^ mice, the number of lesional macrophages did not differ between atherosclerotic *Olfr2*^+/+^ and *Olfr2*^-/-^ mice as previously described by us^19^. Thus, the effect of Olfr2-dependent augmented monocyte recruitment appears to be an effect unique to the AAA model used here.

To this day, AAA remains a highly prevalent disease with little option for intervention. While current treatment focuses on open surgical or endovascular replacement of the aorta, pharmacological targets that may help in preventing or slowing disease progression are lacking. Unfortunately, most clinical trials targeting immune cells and their mediators have been unsuccessful or had to be terminated^52^. Current studies aim for better biomarker profiles (NCT03703947) and investigate CCR2 as an inflammatory marker for PET/CT imaging (NCT04586452). As extra-nasal olfactory receptors only very recently emerged as key players in cardiovascular disease, specific drugs beyond citral targeting Olfr2 are lacking and require extensive development and optimization.

This study has several limitations. As a reduced incidence and AAA development has been reported for female compared to male mice^53, 54^, we here report only on male mice. This sexual dimorphism is consistent with the incidence in human patients^55^. We think, however, that Olfr2 might play a similar role in AAA of female mice and women. While we could link the CX3CR1-CX3CL1 axis as one driver of impaired migration of *Olfr2*^-/-^ monocytes *in vitro* and *in vivo*, further chemokine receptors or adhesion molecules might contribute to the observed phenotype *in vivo*. The levels of octanal were unchanged in the course of AAA. Multiple sources and mechanisms are involved in the generation of local and systemic octanal. We previously reported on lipid peroxidation of lipid macromolecules, which could act as a potential source of octanal^19^. Lipid peroxidation has been identified to also exert a key role in AAA formation^56^. However, the source of octanal in the context of AAA is not clear and its identification will be part of future studies.

In conclusion, we have identified Olfr2 as a critical driver of AAA by promoting monocyte recruitment and subsequent vascular inflammation. Pharmacological inhibition of Olfr2 may represent a promising strategy to prevent AAA progression in the future.

## Acknowledgments

We would like to thank Nadja Klein, Simon Grimm, Christina Vosen, and Katharina Tinaz for their excellent technical support throughout the duration of this research project. We also would like to thank the CECAD Imaging Facility for their superb support for the imaging experiments. Parts of figures were created with BioRender.com.

## Sources of Funding

This work was supported by the Deutsche Forschungsgemeinschaft (GRK 2407: 360043781); SFB TRR259 (397484323 projects A04, A05, A07, A09, C03 and project 535107899); the Center for Molecular Medicine Cologne; the Neven-DuMont Foundation all to Holger Winkels. DFG-INST 216/741-1 FUGB for the Leica Microsystems TCS SP8 Multiphoton; DFG-INST 216/1068-1FUGG for the Zeiss LSM 980 Airyscan2; INST 216/1063-1 FUGG for the Sony MA900 sorter.

## Disclosures

The authors declare no conflict of interest.

## Notes

### Competing Interest Statement

The authors have declared no competing interest.

